# Climate associated genetic variation in *Fagus sylvatica* and potential responses to climate change in the French Alps

**DOI:** 10.1101/849406

**Authors:** Thibaut Capblancq, Xavier Morin, Maya Gueguen, Julien Renaud, Stéphane Lobreaux, Eric Bazin

## Abstract

Local adaptation patterns have been found in many plants and animals, highlighting the genetic heterogeneity of species along their range of distribution. In the next decades, global warming must induce a change in the selective pressures that drive this adaptive variation, forcing a reshuffling of the underlying adaptive allele distributions. For species with low dispersion capacity and long generation time such as trees, the rapidity of the change could imped the migration of beneficial alleles and lower their capacity to track the changing environment. Identifying the main selective pressures driving the adaptive genetic variation is thus necessary when investigating species capacity to respond to global warming. In this study, we investigate the adaptive landscape of *Fagus sylvatica* along a gradient of populations in the French Alps. Using a ddRAD-seq approach, we identified 7,000 SNPs from 570 individuals across 36 different sites. An RDA-derived method allowed us to identify several SNPs that were strongly associated with climatic gradients; moreover, we defined the primary selective gradients along the natural populations of *F. sylvatica* in the Alps. Strong effects of elevation and humidity, which contrast north-western and south-eastern site, were found and were believed to be important drivers of genetic adaptation. Finally, simulations of future genetic landscapes that used these findings predicted a severe range contraction and a shift towards higher altitudes for *F. sylvatica* in the Alps and allowed to identify populations at risk, which could be helpful for future management plans.

## INTRODUCTION

The long-term survival and the distribution of species are triggered by their ability to grow and reproduce in a given set of environmental conditions (Hutchinson 1957). When populations of the same species experience different environments, adaptation to the local conditions may occur. As a result, individuals have a better fitness in their local environment than individuals from other populations (Rehfeldt et al. 2002). Patterns of local adaptation are commonly studied to better understand adaptation and especially the equilibrium between natural selection and gene flow within a set of intra-specific connected populations (Kawecki and Ebert 2004). Better assessing the genetic selective pressure encompassed by the different populations of a species leads to a better understanding of the ecology and distribution of the species and can help refine our vision of species homogeneity over space and time (Poncet *et al*. 2010, Manel *et al*. 2012, Alberto *et al*. 2013a, De Kort *et al*. 2013, Fitzpatrick and Keller 2015).

Recent projections predict major environmental changes by the end of the century (IPCC, 2007 and 2013), and these changes are already impacting biodiversity (Thuiller *et al*. 2005, Bellard *et al*. 2012). This prediction is especially true for mountainous plant and tree species (Pauli *et al*. 2012, Thuiller *et al*. 2014, Duputié *et al*. 2015, Trumbore *et al*. 2015), which are expected to undergo a shift towards higher elevations (Penuelas *et al*. 2003, Walther *et al*. 2005, Parmesan 2006, Beckage *et al*. 2008, Lenoir 2008, Shaw and Etterson 2012). When projections of species’ responses to climate change have been investigated, the studies have frequently been conducted using presence/absence with habitat suitability modelling (Bakkenes *et al*. 2002, Thuiller *et al*. 2005). In such models, in most cases, the species has been considered as a homogeneous unit, and any potential heterogeneous genetic adaptations across populations have been neglected (Alberto *et al*. 2013a, Bay et al. 2017a). Nonetheless, in the last decade, it has been proposed that genetic heterogeneity should be considered when studying species diversity and distribution (Jay *et al*. 2012, Alsos *et al*. 2012, Steane *et al*. 2014, Razgour *et al*. 2018). To do so, the goal would be to translate the genomic information into spatially explicit predictions of genetic and especially adaptive variation (Schoville et al. 2012, Fitzpatrick and Keller 2015). Furthermore, some authors have suggested that local adaptation must be integrated into estimates of the risk of species’ range losses under climate change scenarios (Morin *et al*. 2007, Benito Garzon *et al*. 2011, Mouquet *et al*. 2015, Bay et al. 2017b, Bay et al. 2018, Exposito-Alonso *et al*. 2018, Martins *et al*. 2018).

The adaptive genetic heterogeneity could largely impact species response to climate change in the next decades (Aitken *et al*. 2008, Jump & Penuelas 2005, Rehfeld *et al*. 2002). A changing environment induces a continuous modification of the selective pressure across the landscape and could create a distortion between population phenotypes and newly optimal ones (Hoffmann & Sgro 2011, Aitken *et al*. 2008). It would result in a need for the populations to change their adaptive genetic component accordingly and track the new optima across the landscape. An emerging field of research aims at estimating the magnitude of the deviance between the current and future optimal genetic composition (Steane *et al*. 2014, Fitzpatrick & Keller 2015, Bay *et al*. 2018, Exposito-Alonso *et al*. 2018, Martins *et al*. 2018). To do so they investigate the relationship among genetic, phenotypic and environmental variability and extrapolate these relations across a landscape. By looking at the deviance between optimal genetic composition in current and future conditions they estimate a potential genetic offset (Fitzpatrick & Keller 2015) that the populations would have to reduce to ensure their survival. These new methods have been facilitated by the recent development of next-generation sequencing (NGS) techniques, which have created the possibility to access a large amount of genetic variation across the genome (da Fonseca *et al*. 2016, Hoban *et al*. 2016).

The present study uses a RDA-based approach and a large matrix of genomic data to investigate the genetic-environment relationship and estimate the potential deviance between current and future genetic composition optima in common beech (*Fagus sylvatica*) along the French Alps. *F. sylvatica* is a tree species commonly found in European mountain ecosystems, covering large environmental gradients, with a distribution from the Pyrenees Mountains to Scandinavia (Frydl *et al*. 2011). In the Alps, the species is distributed at altitudes ranging from 600 to 1800 m. This distribution spans a variety of environmental conditions, sometimes at very small geographical scales (e.g., along altitudinal gradients), which makes beech populations likely to be affected by local adaptation. A recent study by Garate-Escamilla *et al*. (2019) identified local adaptation to variation in potential evapotranspiration in beech populations across Europe. Furthermore, two other studies by Csilléry *et al*. (2014) and Pluess *et al*. (2016) found evidence for local adaptation to climate, as the authors identified adaptive variation at very short spatial scales (< 10 km) in the south of the French Alps and at a regional scale across Switzerland (> 100 km), respectively. Thus, beech appears to be an appropriate species to identify the adaptive component of genetic variability, determine the environmental factors shaping this component variation in populations, and explore the potential impact of climate change on the future adaptive landscape of the species.

Here, we aimed to i) identify the environmental constraints partially shaping the genetic differentiation of beech along the French Alps; ii) isolate the genetic variation associated with environmental gradients; and iii) consider genetic heterogeneity to estimate the species’ capacity to respond to climate change. To answer these questions, we based our investigations on a single-nucleotide polymorphism (SNP) dataset obtained through a double-digest restriction-site associated DNA sequencing procedure (ddRADseq, Peterson *et al*. 2012). We used a large number of SNPs genotyped in our sampling to combine analyses of genetic differentiation, genetic variance decomposition and genome scans. This approach allowed us to identify the main climatic drivers of *F. sylvatica* genetic variation and to extrapolate the contemporary and future adaptive potential of this species along the French Alps.

## MATERIALS AND METHODS

### 1. Material and genetic data acquisition

We sampled 36 populations of *F. sylvatica* along the western front of the French Alps (Fig. 1 and Table S1) during the summer of 2016. The complete sampling included 570 individuals distributed in 19 different mountain ranges located from the Jura to the Mediterranean Alps. This sampling covered most of the distribution of *Fagus sylvatica* in the French Alps.

**Figure 1:**
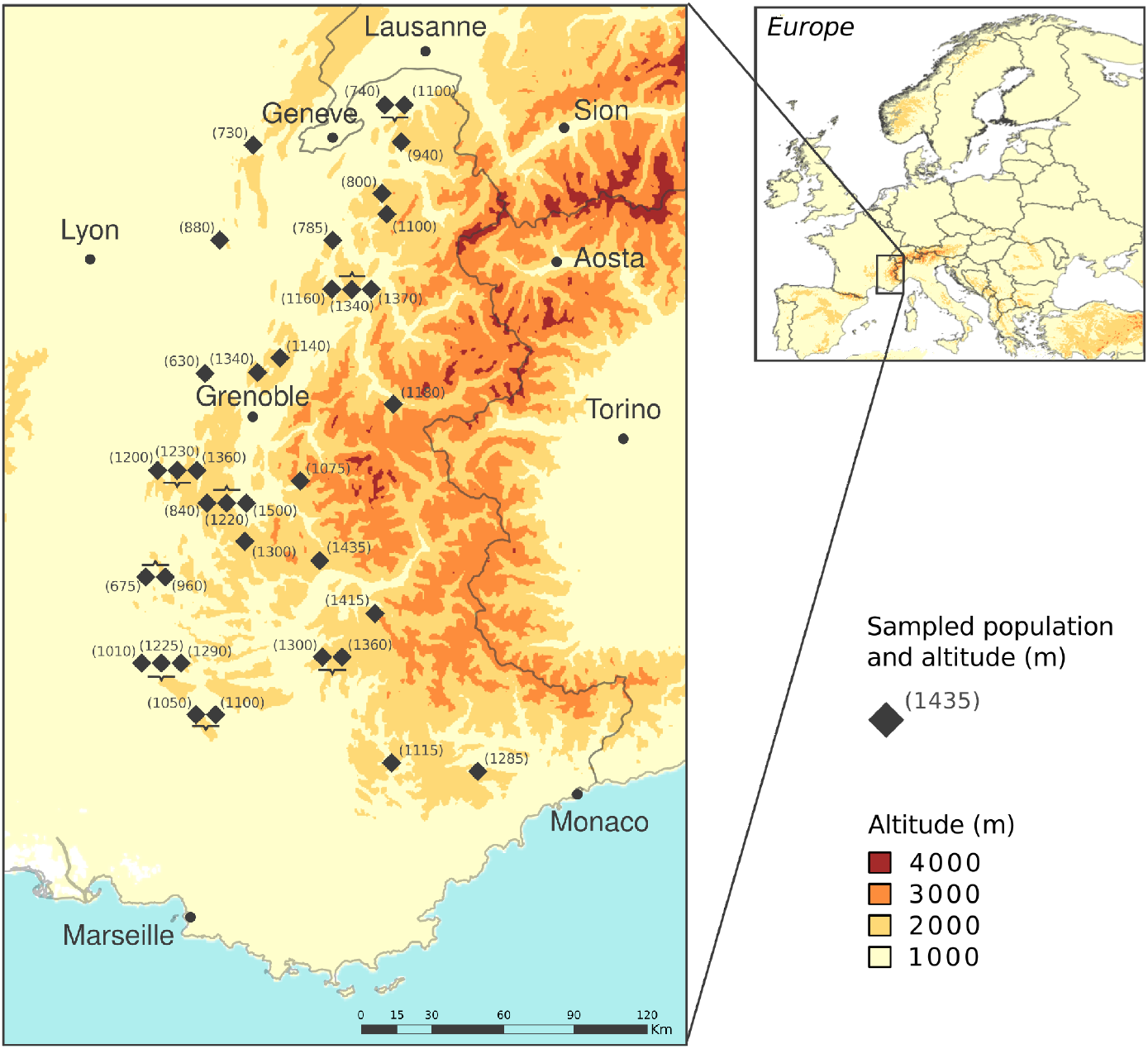
Map of the study area with locations of sampled population. The numbers give the number of sampled individuals in each population.

#### ddRADseq protocol

DNA was extracted from one or two leaves of each individual using the DNeasy Plant kit (QIAgen) according to the manufacturer’s instructions, and samples were stored at −20°C. A double-digested RAD (restriction site associated DNA) experiment was conducted on the 570 individuals (using 12 different libraries) using a modified version of the protocol described in Peterson *et al*. (2012). The enzymes used were PstI and MspI; for more details on the experimental protocol, see Capblancq et al. (2015). The 12 libraries were then sequenced on a complete lane of Illumina Hi-Seq 2500 2x 125 (Fasteris SA, Switzerland). DNA reads (∼250 million reads) resulting from the sequencing were used to genotype single-nucleotide polymorphism (SNP).

#### Sequences treatment

We demultiplexed the sequences with the program *STACKS* (Catchen *et al*. 2013) using a Phred score of 10 for reads filtering (*process_radtags* function). We mapped the reads against the newly assembled beech genome (Mishra *et al*. 2018) using BWA software (Li and Durbin 2009) and the BWA-MEM algorithm. We processed the mapped sequences with the *gstacks* function of *STACKS* pipeline using a minimum mapping quality of 20. Finally, only SNPs on tags present in at least 60% of the 570 individuals and with a frequency higher than 0.5% (∼3 individuals) were used for further analyses.

Using the *ProcessMyRAD* scripts (https://github.com/cumtr/PmR, Cumer *et al*. 2018), we also produced graphical outputs at the filtering step of the treatment, which allowed an evaluation of the experimental success for each library and individual. This method provides the proportion of retained reads after the first step of filtering, the number of reads by individuals, the quality of the sequencing and the proportion of the different nucleotides along the reads.

### 2. Genetic variation among the populations

We estimated individual grouping and population differentiation using principal component analysis (PCA) that was conducted on the ddRADseq SNPs dataset using the *adegenet* R package (Jombart 2008). We also estimated the population structure using a constrained method of genetic clustering: sNMF (Frichot *et al*. 2014). This analysis was performed using the R package *LEA* (Frichot and François 2015), with the number of genetic clusters (K) ranging from 2 to 6.

### 3. Testing genetic-environment relationship

To detect genetic-environment association in alpine populations of *Fagus sylvatica*, we tested the link between a set of climatic variables and the genetic variability across the sampled populations. Specifically, we used a series of partial redundancy analysis (RDA) performed using the function *rda* of the *vegan* package in R (Oksanen *et al*. 2015). The genetic dataset (e.g., allelic frequencies in populations in our case) was used as response matrix Y, and a set of environmental variables was used as explanatory matrix X. A third matrix of geographic variables and “neutral” genetic composition was used as a conditioning matrix to avoid confounding associations between genetic variability and geography and/or evolutionary history. This conditioning matrix allowed the consideration of a potential pattern of isolation caused by distance or a genetic differentiation due to lineage splitting rather than adaptation to the environment. All the different combinations of the explanatory and conditioning variables have been used in the RDA model to partition the percentage of genetic variance explained by each specific set of variables (e.g., climatic, geography or “global/neutral” genetic composition).

The geographic variables used in the partial RDA models are typically the coordinates of the population (longitude: x and latitude: y), and the “neutral” genetic composition is the ancestry coefficient obtained from the sNMF analysis for K=2 (see section *Genetic variation among the populations*). We used longitude and latitude without any correction for the curved surface of the planet because of the small geographic scale considered here. The set of climatic variables used in this study came from Thuiller *et al*. (2014), and these variables are known to be drivers of plant distribution over an area, such as the French Alps. These variables include the annual sum of degree-days above 0°C (*degg0*), the annual mean potential evapotranspiration (*etp_mean*), a moisture index (*mind_mean*), the sum of the annual precipitation (*prec_sum*), the maximum temperature (*tmax_mean*) and the minimum temperature (*tmin_mean*). These variables were available at 250m-resolution in the studied area (see Thuiller *et al*. 2014 for more details). To avoid multi-collinearity between variables, only variables that were not overly correlated (i.e., R-squared < 0.8) were included in the final analysis, which avoids overfitting due to redundant information (Fig. S2).

### 4. Signature of adaptation

Specific loci associated with environmental adaptation were searched using a method derived from RDA and following the procedure proposed in Capblancq *et al*. (2018). The approach uses RDA to identify loci that are extremely linked to environmental variables and are likely under selection in the sampled population (see Lasky *et al*. 2012, Forester *et al*. 2016 & 2017, Capblancq *et al*. 2018). This method starts using a classical RDA procedure, and an outlier locus is detected when its projection in the K first axis of the RDA (principal component) does not follow the projection of the majority of the loci along the same K principal component. More precisely, outliers correspond to loci for which the Mahalanobis distance (i.e., the distance estimated between the z-scores of the locus on the K first principal component and the mean of all the loci z-scores on the same set of principal components) is extreme relative to the distribution of the Mahalanobis distances of all loci (Luu *et al*. 2017, Capblancq *et al*. 2018). A p-value is then obtained for each locus after a transformation of the Mahalanobis distances (see the procedure in Luu et al. (2017)).

In this study, we used the population allelic frequencies as the genetic matrix (response matrix Y) rather than individual genotypes; this avoided biases due to a non-equal number of samples in the genotyped populations. Only the SNP with a minor allele frequency superior to 10% of the complete sampling (> 5 indidivuals). Furthermore, we wanted to consider the past evolutionary history of the population during the RDA, and thus, we conditioned the analysis using the global genetic structure observed in the sampling. To do so, we used the ancestry coefficient obtained from the sNMF analysis for K=2 (see section *Genetic variation among the populations*). We assumed here that the ancestry is a pertinent proxy to catch the neutral genetic differentiation due to both isolation caused by distance among populations and demographic history of the populations (*i.e.* post glacial colonization history). The use of ancestry as a conditioning variable could remove any variation linked to both geography and environment. We resolved this issue by sampling multiple populations showing strong environmental differentiations (*i.e.* differences in altitude, see Figure 1) at very short distance. Finally, we controlled for a false discovery rate (FDR) by transforming the p-values into q-values using the procedure of the *qvalue* R package (Storey, 2002). We kept the loci with q-values less than 10^-4^, which corresponds to a false discovery rate of 0.01%.

### 5. Spatial extrapolation of the adaptive landscape

#### Adaptive indices

We performed a second RDA with only the loci previously found as outliers (*q.values* < 0.0001). This set of outliers provides an “adaptively enriched genetic space” (Steane *et al*. 2014), and a second RDA on these specific loci allows the identification of environmental variables that are the most correlated with putative adaptive variation. Thus, we used the scores of the different environmental variables along the first two RDA axes to build composite indices that predicted the adaptive score of individuals in the environment following the formula:

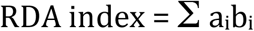

where *a* is the variable score (loading) along the concerned RDA axis, *b* is the value of the standardized environmental variable in the concerned location, and *i* refers to the different variables used in the RDA model (Steane *et al*. 2014). Adaptive indices were estimated in the entire study area for both RDA1 and RDA2. A map representation of these indices allows for a better understanding of the adaptive landscape of *F. sylvatica* in the French Alps.

#### Future predictions

The same climatic variables previously used were projected for 2050-2080 following the A1B emission scenario (see Thuiller *et al*. 2014 for more details). In the same way as it was conducted for the current climatic conditions, we used these projected variables to spatially predict the RDA1 and RDA2 indices in the future. We then used the present occurrences of *F. sylvatica* in the French Alps (data came from the French National Alpine Botanic Conservatory, CBNA, and the National Mediterranean Botanic Conservatory, CBNMED) to extract the current favourable range of the RDA1 and RDA2 indices and to compare them to the indices values predicted in 2080 using the A1B scenario.

#### Assessing the capacity of species’ responses to climate change

For each pixel included in the current range of *Fagus sylvatica* in the French Alps, we estimated a need of adaptive score change, equivalent to the genetic offset proposed by Fitzpatrick and Keller (2015). To do so, we subtracted for both RDA1 and RDA2 the value of the index predicted with the present climatic conditions and the value of the index predicted with the future climatic conditions. We thus obtained a specific genetic offset for each of the two RDA axis and sum them to obtain an estimate of the global genetic offset. Moreover, for each future favourable pixel, we measured the distance to the closest population showing an equivalent adaptive score under the current climatic conditions. For this purpose, we looked for both RDA1 and RDA2 the closest pixel showing an index being in the same quartile under the current climatic conditions and averaged for each pixel the two distances obtained (RDA1 and RDA2). With this index we wanted to estimate a resistance of adaptation potentially occurring if the alleles favourable in the future climatic conditions have to migrate long distances to reach the focal location. Finally, we investigate the levels of adaptive standing genetic variation (SGV) in the populations by calculating the mean of the outlier loci allele frequency variances (*p x q*), as an index of within population adaptive variation (Chhatre *et al*. 2019). In the same way, we estimated the Population Adaptive Index (PAI) of each population as proposed by Bonin et al. (2007). PAI is calculated by measuring the absolute difference between the adaptive allele frequencies of a specific population and the mean adaptive allele frequencies of the complete sampling. When the estimation of allele frequency variance gives a measure of the availability of the adaptive alleles in the population (Standing Genetic Variation), the PAI gives an estimate of the extremeness of the population along the gradient of genetic adaptation.

## RESULTS

### 1. *Fagus sylvatica* genetic variability along the French Alps

The double-digested RAD libraries produced a mean of 11,588 fragments, with a mean coverage of 22.6 reads/fragment for the 569 samples that were analysed. The polymorphism in the RAD fragments allowed the scoring of 7,010 independent SNPs with 19% of missing data at the end. PCA on these SNPs succeeded in differentiating the sampled populations of *Fagus sylvatica*. The differentiation was mostly carried by the first PC axis, which exhibited a clear north-south differentiation pattern with high scores for the Mediterranean populations (e.g., Prealpes-Azur or Verdon) and negative scores for the northern populations (Fig. 2). These results were congruent with the results of genetic clustering analysis using sNMF. The sNMF analysis showed a progressive gradient of assignation when two clusters were constrained with the same north-south pattern of differentiation (Fig. 2). When looking at higher values of K, we can see that new cluster assignations did not identify clear genetic groups of individuals and/or were not really consistent with geography (Fig. S3).

**Figure 2:**
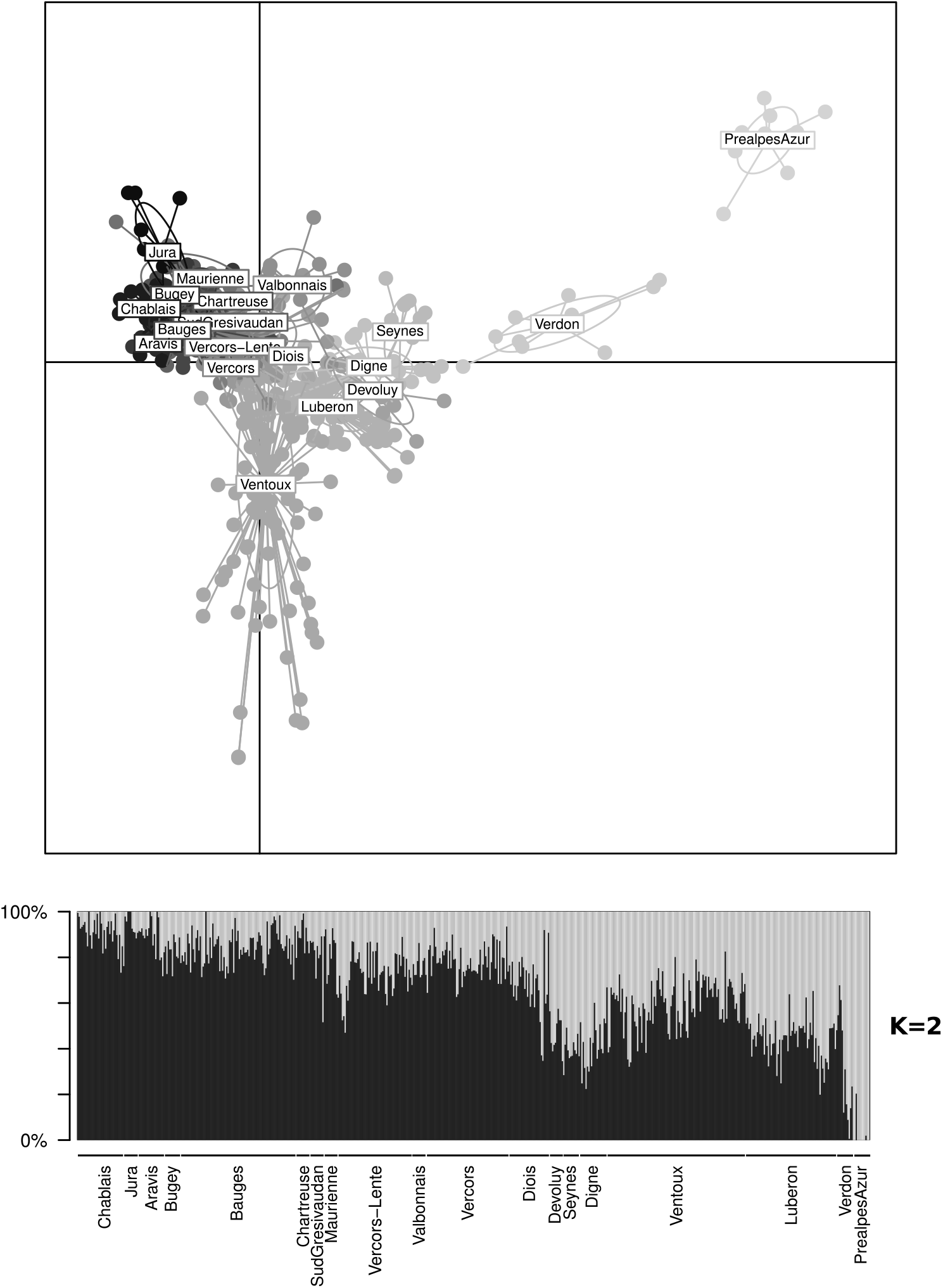
Principal component analysis (top) and sNMF clustering (bottom) obtained from the ddRADseq genetic data set. On the PCA plot, the first two PCs are represented; they caught 2% of the total variance (1.16% for PC1 and 0.81% for PC2). The ellipse gathered the different populations and the colour of the points and ellipses are linked to their latitude with black points coming from northern populations and grey points coming from the populations close to the Mediterranean Sea. The sNMF results are shown for K = 2, the samples are ordered depending on their latitudinal coordinates with northern locations in the left of the plot.

### 2. Genetic-environment association

The different partial RDAs performed on the population allelic frequencies allowed the identification of the proportion of genetic variance independently explained by the climatic variables, geography or ancestry of the populations. The climatic variables used in the analysis included the mean evapotranspiration (*etp_mean*), the maximum and minimum temperature (*tmax_mean, tmin_mean*), the sum of the annual precipitation (*prec_sum*) and the moisture index (*mind_mean*). We removed the other variables because they were strongly correlated with the previous ones (Fig. S4). For all the RDAs and to avoid potential correlation bias due to rare values, we removed all the loci with minor allele frequencies inferior to 1% of the sampling, leaving us with 6,857 loci for which allele frequencies were calculated for all of the populations.

The model that considered all the variables (e.g., climatic variables, geographic variation and ancestry) produced a strong significant association between these variables and the allelic frequencies in the populations. This model explained 35.2% of the total genetic variability (Table 1). The different partial RDAs identified that 47.1% of this global explained variance was associated with climatic variation only (16.6% of the total genetic variance), 19.4% was due to geographic variations (geographic coordinates), 13.9% was due to the ancestry of the individuals and 19.6% was not distinguishable among these different parameters. The five climatic variables used were able to explain a considerable and significant part of the genetic variability, suggesting that environmental conditions have constrained a portion of the genetic composition in *F. sylvatica* populations along the Alps.

**Table 1:** Table of partial redundancy analysis results. The influences of each set of variables (climate, geography and ancestry variables) have been tested separately and together. The percentage of explained total genetic variance is given for each of the model together with the significance of the test and the percentage of variance explainable by the complete set of variables.

|  | Inertia | p(>F) | Proportion of total<br>Variance |
| --- | --- | --- | --- |
| **Total explainable : $F \sim clim. + anc. + geog.$** | **21.92** | **0.001 \*\*\*** | **0.333** |
| **Pure climate : $F \sim clim. \mid (geog. + anc.)$** | **11.136** | **0.001 \*\*\*** | **0.169** |
| **Pure geography : $F \sim geog. \mid (clim. + anc.)$** | **6.04** | **0.001 \*\*\*** | **0.092** |
| **Pure ancestry : $F \sim anc. \mid (clim. + geog.)$** | **1.95** | **0.001 \*\*\*** | **0.03** |
| **Confounded climate/geography/ancestry** | **2.794** |  | **0.043** |
| **Total unexplained** | **43.82** |  | **0.667** |
| **Total inertie** | **65.74** |  | **1** |

Building on these results, we conducted the RDA-based genome scan procedure using the same five non-correlated climatic variables and the ancestry of the individuals for *K =* 2 (averaged by population), as a conditioning matrix. We retained the 3 first axes to process the genome scan, as indicated by the screeplots (Fig. S4). The analysis with an FDR of 10^-4^ retains 65 loci as outliers among the 6,857 tested SNPs (Fig. 3A). The part of the genetic sampling represented by these 65 loci (0.01% of the SNPs) showed a strong association with environmental variation and was hypothesized to represent genomic regions associated with environmental selection in *F. sylvatica*. This set of 65 highly significant loci was then considered as the “adaptively enriched genetic space” of *Fagus sylvatica* in the Alps. They served as a base to extrapolate the adaptive constraints encompassed by the species in the sampling area and to analyse the adaptive landscape.

**Figure 3:**
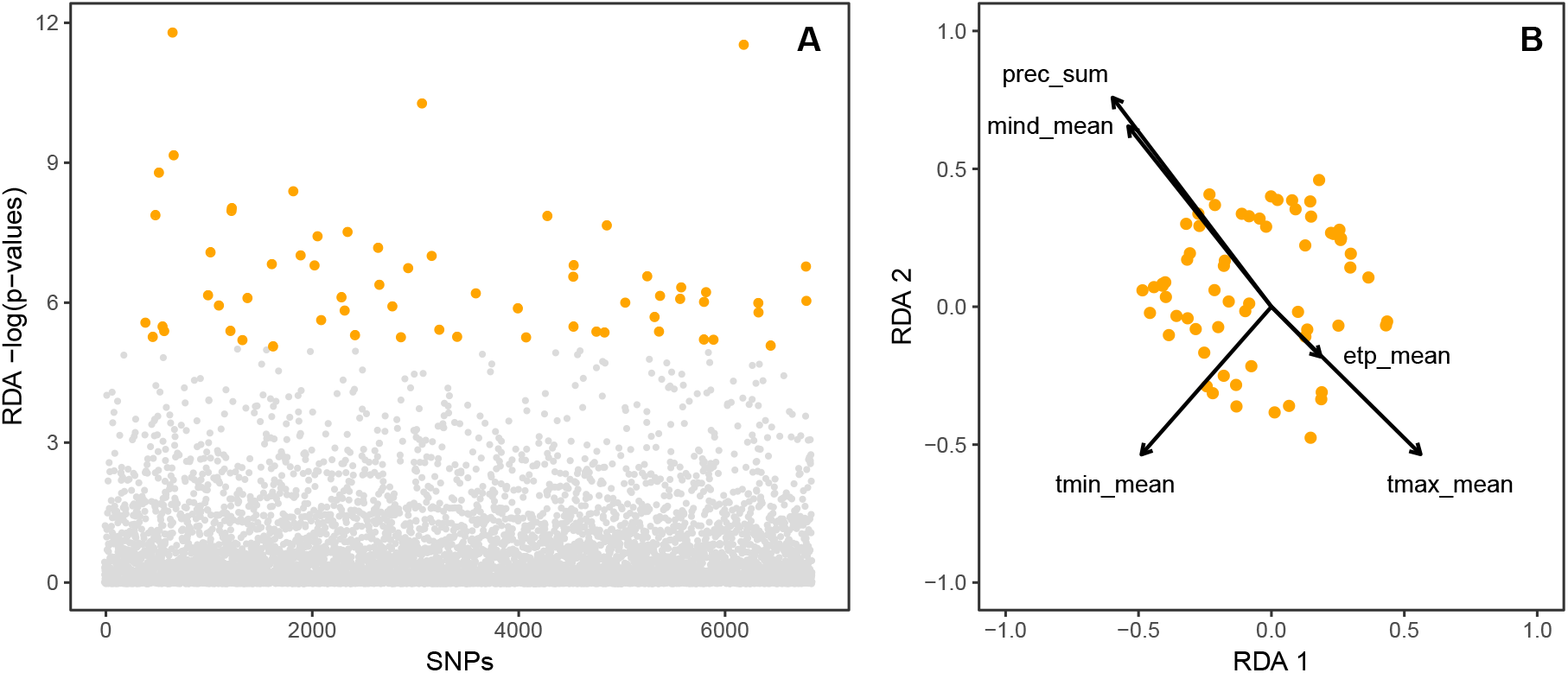
(**A**) Manhattan plots of the RDA results. Loci with a q-value superior to 0.1 % are colored in orange. The ddRADseq loci are not localised in the genome, in this way the x-axis is not informative on loci proximities. (**B**) Projection of loci and environmental variables into the adaptively enriched genetic RDA space. The two first axes are represented; they explain respectively 36% and 20% of the total variance.

### 3. Adaptive landscape

When we performed a new RDA on the 65 outlier SNPs constituting the adaptively enriched genetic space, we found that the two first axes explained most of the adaptive genetic variance among the populations (*i.e.*, 35 and 20%, respectively). RDA1 was correlated with most of the environmental variables used in the analysis and contrasted high maximum temperatures from high values of minimum temperature, precipitation and moisture index (Fig. 3B). Such gradient contrasts the populations from northern locations in the pre-alps where the minimum temperatures are not very low and the precipitation are important with the southern mountainous populations where the minimum temperature is usually lower but with a drier climate. In the other side, the RDA2 was correlated with another part of the variation that contrasted high values of maximum and minimum temperature to high values of precipitation or moisture index. Thus, RDA2 was clearly associated with the altitudinal constraints.

In the same way, the correlation between the different outlier SNPs and the RDA axes showed interesting patterns (Fig. 3A and Fig. S5). Some of the 65 outliers were specifically associated with one of the two RDA axes, demonstrating that the genetic variation associated with RDA1 and RDA2 was partially supported by different loci (Fig. S5). The RDA procedure orientated ordination axes in the direction of orthogonal environmental gradients. The two observable sets of adaptive loci were thus “independently” associated with the two different environmental constraints experienced by *F. sylvatica* in the French Alps. Nonetheless, many loci were associated with both RDA1 and RDA2. Finally, the variation in the allelic frequencies across the different sampled locations was high for all 65 outlier SNPs (Fig. S5). The frequency differential varied from at least 0.50% and up to 0.8%, showing that outlier identification was not based only on small variations.

We extrapolated the adaptive genetic variation to the entire French Alps by estimating the RDA1 and RDA2 scores in the entire range of *F. sylvatica* in the French Alps (Fig. 4). The two gradients showed some similarities but the RDA1 index was mainly differentiating the mountain ranges receiving more precipitation in the northern Alps (i.e., Chartreuse, Bauges, Aravis, Chablais, Jura) from the rest of the study area, especially the valleys (i.e., Durance valley, Haute-Maurienne) and the hinterland of the Provence region (Fig. 4) that are typically much drier. On the other side, the RDA2 index was mostly impacted by the altitude of the location. The highest values of this index corresponded to high elevations, and the lowest values were found near the Mediterranean Sea or in lowlands along the Rhone River. 99% of the current occurrences of *F. sylvatica* had an RDA2 index value ranging between −0.93 and 5.99 (Fig. 5). These two values probably represented the limits of the favourable range for *F. sylvatica* along this adaptive constraint.

**Figure 4:**
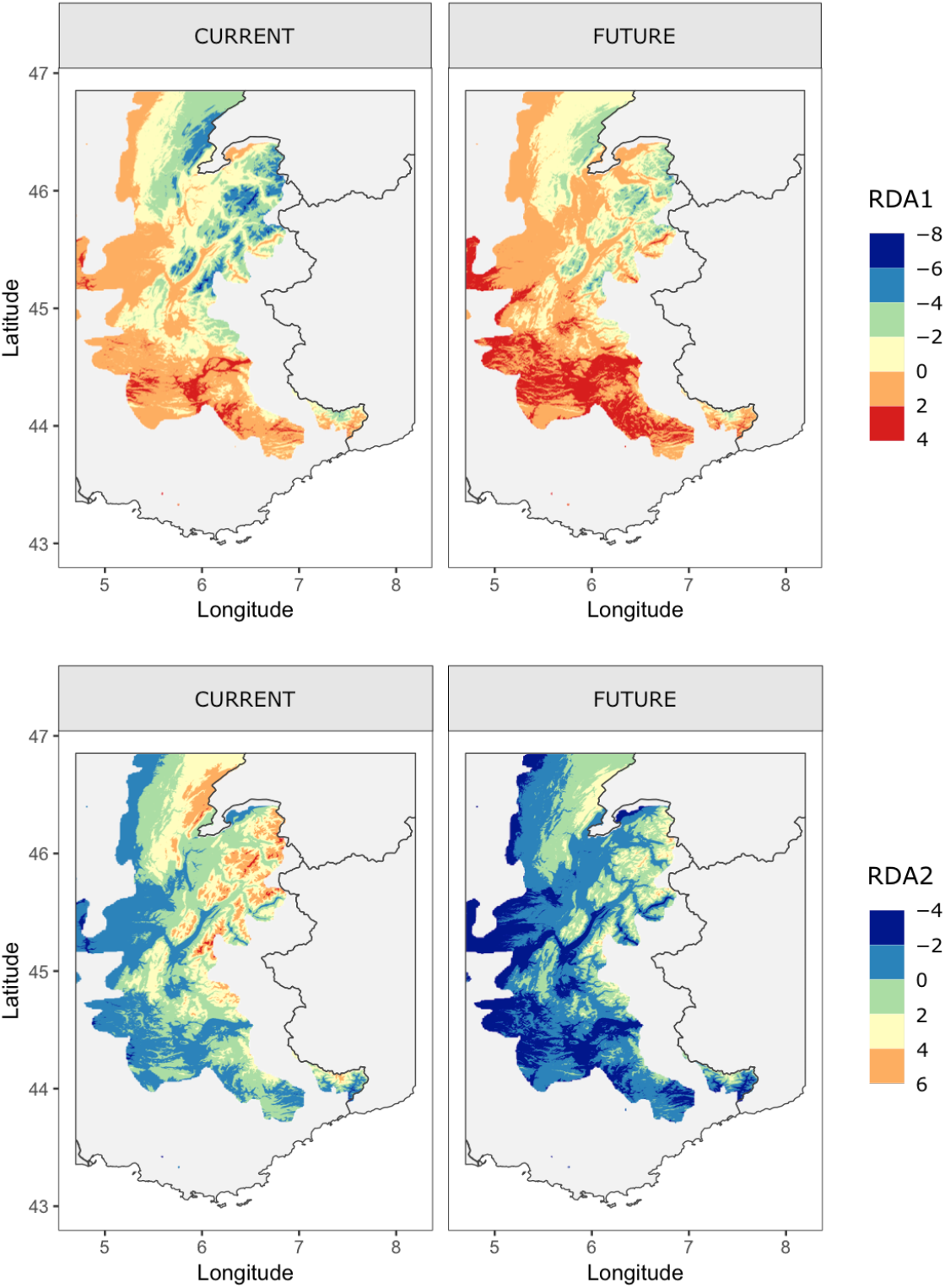
Spatial extrapolation of RDA1 and RDA2 gradients in the French Alps for the current climatic conditions and with future climatic conditions predicted by A1B scenario. The bottom panels show the impact of climate change in *Fagus sylvatica* genetic landscape along the French Alps.

**Figure 5:**
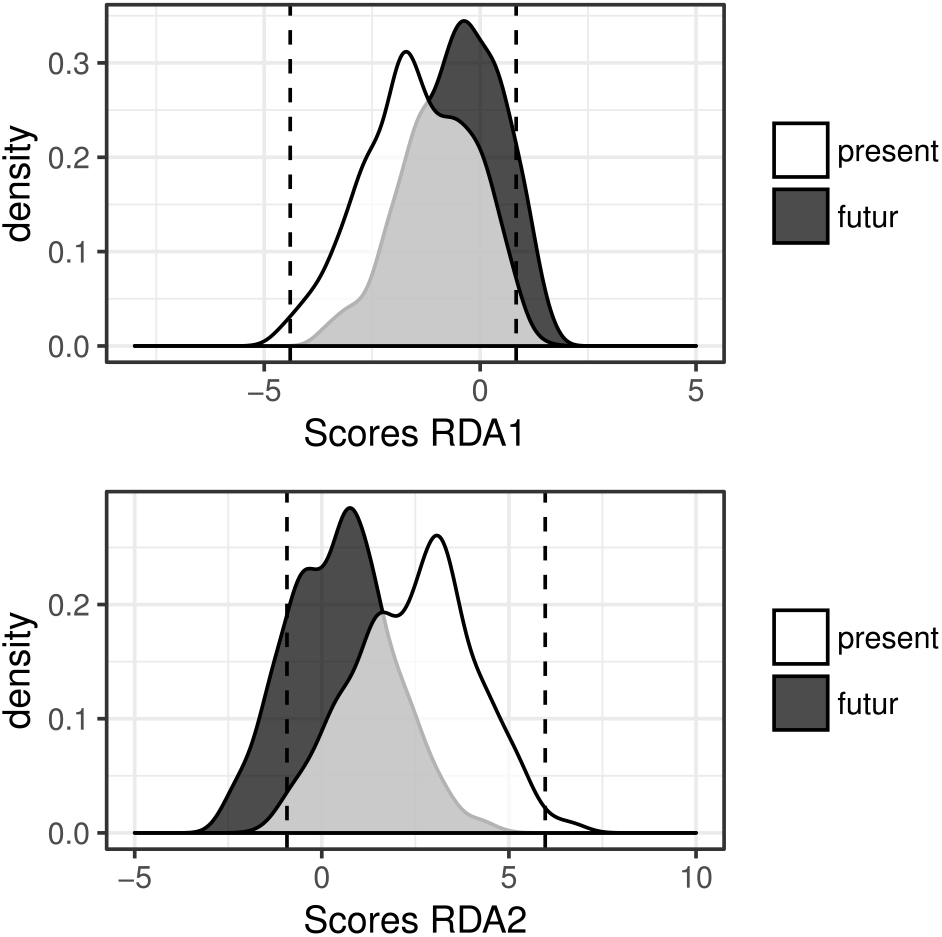
Distribution of RDA1 and RDA2 scores in present occurrences of *Fagus sylvatica* in the French Alps. The distribution of RAD scores is given with actual climatic conditions (top) and future estimated climatic values (bottom). The dashed lines represent the 95% interval of present RDA scores distribution. The black points indicate the actual scores of the sampled populations.

### 4. Potential capacity of response to climate change

Regarding the future predictions made for 2080 using the A1B scenario of climate change, we found a change in the RDA1 index would especially concern the southern areas of the range where the future index values (pixels with a value >2 in Fig. 4) would exceed the values observed for the current climatic conditions (Fig. 4 and Fig. 5). However, we found that the predicted change of the RDA2 index was even more dramatic. According to the predictions, almost one-quarter of the current location of *F. sylvatica* would be beyond the RDA2 index favourable range in 2080 (Fig. 5). In this case, the change would mainly affect the lowland parts of the range where the RDA2 values would go under the lower limit of the index in the current climatic conditions (values < −2 in Fig. 4 and Fig. 5).

Different proxies have been measured to investigate the capacity of *F. sylvatica* to respond to climate change across the French Alps. The need of adaptive score change between the current and future climatic conditions (Fig. 6A) and the distance of populations showing current adaptive scores equivalent to the future adaptive needs (Fig. 6B) have been estimated by combining the RDA1 and RDA2 indexes (see Materials and Methods). These two proxies, respectively called genetic and geographic offset, have been estimated across the current range of *F. sylvatica* in the French Alps, not taking into account potential future colonization of newly favourable locations. We first observed that none of them were homogeneous across the studied area, the genetic offset being greater in mountains areas (Fig. 6A), while the highest values for the geographic offset were found in the valleys and at low elevations (Fig. 6B).

**Figure 6:**
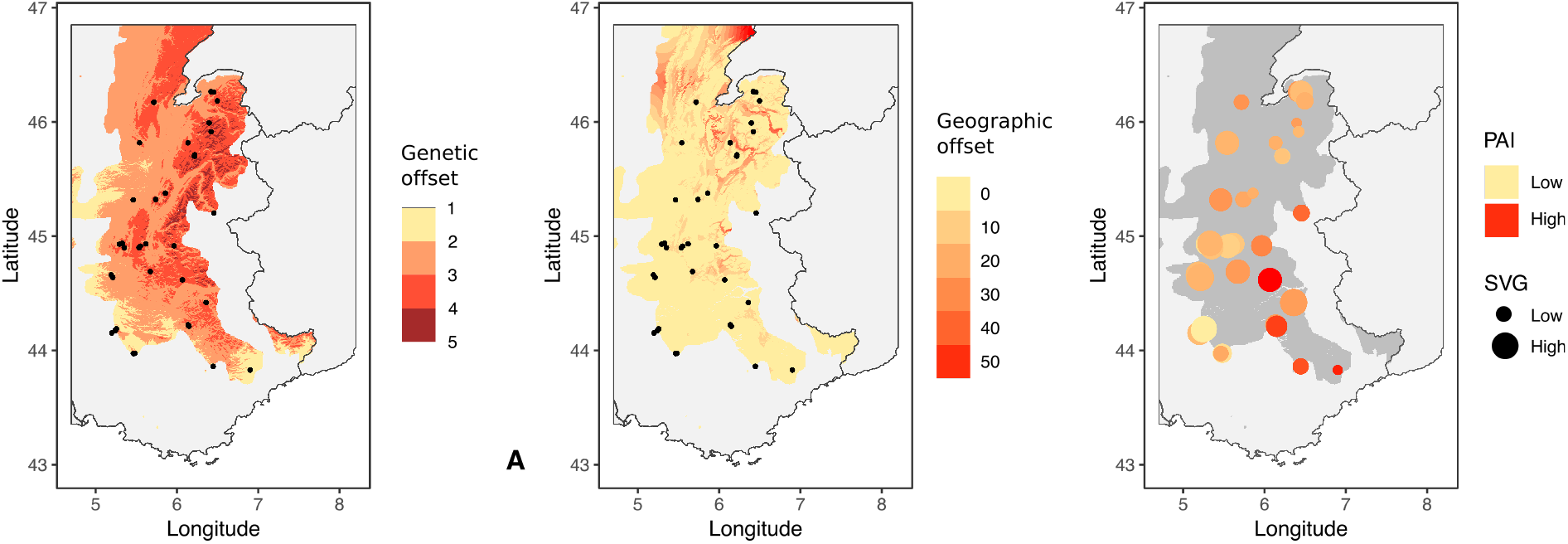
Spatial representation, in the future favourable locations for *F. sylvatica,* of (A) RDA1 index difference between current and future climatic conditions, (B) distances from populations currently showing a RDA1 score equivalent to the future adaptive needs and (C) a composite index taking the two previous factors into account.

Finally, when looking at the actual adaptive genetic variation in the sampled populations, we found that the locations sampled in the southern part of the Alps showed a combination of low rate of standing genetic variation and high value of PAI (Fig. 6C). The six populations sampled in the Chartreuse and Bauges mountains also showed weak level of SGV, but a relatively low PAI. One population sampled at relatively high elevation in the Devoluy mountains showed, at the opposite, a combination of high SGV and PAI.

## DISCUSSION

### Fagus sylvatica genetic variability and evidence of association between genetic and environmental variation

Benefitting from the large genetic matrix obtained through ddRADseq sequencing, we found a substantial genetic variability across the *Fagus sylvatica* populations of the French Alps. We observed a gradual genetic differentiation between the northern locations (i.e., Jura or Chablais mountain ranges) and the locations close to the Mediterranean Sea (Fig. 2). This latitudinal gradient of genetic composition suggests a pattern of isolation by distance without a clear geographical barrier to gene flow among *F. sylvatica* populations. The absence of discrete genetic groups is also congruent with the history of postglacial colonization proposed for *F. sylvatica* by Magri and colleagues (2006) and by Magri (2008). In these studies, the authors suggested that the north-western French Alps would have served as a refuge for *Fagus sylvatica* during the Last Glacial Maximum (LGM). The species would have colonized all the French Alps from this particular location since the end of the glacial period.

Disentangling the signatures of isolation by distance (IBD), local adaptation and demographic history in the genetic variation of a species can be tricky (Nadeau et al. 2016). IBD is largely known to induce neutral genetic gradients that are strongly correlated to the distance among populations (Meirmans 2012, Orsini *et al*. 2013), and postglacial colonization is able to produce allele frequency gradients similar to IBD as a result of repeated founder events along the colonization front (De Lafontaine et al. 2013). Thus, it is difficult to separate the effects of geography, environment and demographic history when they are spatially correlated, as seems to be the case for *F. sylvatica* in the French Alps. The use of partial RDAs allowed us to tackle this issue by decomposing the portion of genetic variance explained by each group of variables when any potential associations with the other variables have been removed (Legendre and Legendre 2012). Our results showed that a proportion of *F. sylvatica* genetic variation was explained by geographic distance (7%), demographic history (5%) and climate (17%). A confounding effect of the three factors was also found (7% of the variation), attesting the difficulty of completely disentangling these variables (Table 1).

A great portion of the genetic variation was explained by climate (17%), which suggests that environmental variables partially shaped the genetic composition of *F. sylvatica* populations across the sampling area. We suggest here that local adaptation occurs in *F. sylvatica* populations along the French Alps, as has already been evidenced in Mount Ventoux (France, Csilléry *et al*. 2014), in Switzerland (Pluess *et al*. 2016) and at very short geographic scales along an altitudinal gradient in two adjacent valleys of the Pyrenees (Vitasse *et al*. 2009). At larger geographic scale, provenance trials across whole Europa have revealed population differentiations suggesting local adaptation (Gömöry & Paule 2011, Kreyling *et al*. 2014) and a recent study investigating at the same time the effects of plasticity and genetically driven adaptation also found evidence of local adaptation to potential evapotranspiration across the whole range of *F. sylvatica* in Europe (Garate-Escamilla *et al*. 2019). Our results are therefore complementary to these previous works, confirming the idea that this species is a good model to further investigate the factors driving species adaptation and distribution.

### The factors driving adaptive genetic variation in F. sylvatica

The statistical approach we used to investigate the drivers of local adaptation in *F. sylvatica* is based on redundancy analysis (RDA) and is just starting to show its potential in the field of genetics of adaptation (Lasky et al. 2012, De Kort *et al*. 2014, Steane *et al*. 2014, Forester *et al*. 2016 and 2017, Capblancq *et al*. 2018). RDA has shown great potential in identifying the signatures of selection and adaptive loci, both in comparison with other genome scan methods (Forester *et al*. 2017, Capblancq *et al*. 2018) and in different types of environments (Forester *et al*. 2016). Such multivariate approaches are thought to be more efficient in detecting multi-loci selection since they consider the potential co-variation among genetic markers (Rellstab *et al*. 2015). This fact is important to highlight because we know that traits involved in local adaptation seem to be triggered by many genes that generally have weak effects (Aitken *et al*. 2008, Savolainen *et al*. 2013, Yeaman 2015, Bay *et al*. 2017a). Moreover, by allowing the identification of the environmental gradients that are associated with adaptive genetic variation, this approach can be used to spatially predict a landscape of species adaptive capacity under current or future environmental conditions (Steane *et al*. 2014, Fitzpatrick and Keller 2015, Capblancq *et al*. 2018).

Here, we found that two principal environmental gradients have triggered two relatively independent adaptive genetic responses in the French alpine populations of *Fagus sylvatica* (Figs. 3 & 4). The first adaptive genetic variation identified was associated with both humidity and minimum and maximum temperatures variations (RDA1 in Fig. 3). This gradient contrasts the northern mountainous locations (*i.e.* high elevation) with the southern and valleys locations (*i.e.* low elevation) of the sampling zone (Fig. 4). Previous studies have described that southern and internal zones (*e.g*. Maurienne valley) of the Alps receive less rainfall that impact the survival of beech populations (Ozenda 1985, Courbaud *et al*. 2011). These regions showed the highest RDA1 index values (Fig. 4), with values very close the extreme edge of the species range in term of survival capacities (Fig. 5). Furthermore, this constraint is not the only one exercising a selective pressure on the *F. sylvatica* genetic variation. We found a second adaptive genetic gradient linked to environmental variations induced by the altitudinal gradient (RDA2 in Fig. 3 & Fig. 4). Such finding strongly suggests that the identified outlier genetic markers are close to genomic regions involved in *F. sylvatica* adaptation to the differential climatic conditions between lowlands and high altitudes. A signature of genetic adaptation correlated with the altitudinal gradient has similarly been found by Csilléry and colleagues (2014) in one population of the south-western French Alps (Mount Ventoux).

To temper our results, it is important to note that, given a genome size of 535 Mbp (Kremer 2012), the 6,857 genotyped SNPs would be spaced at about 78, 000 bp on average. Linkage disequilibrium probably decays much more rapidly, even around selected sites. We are thus aware that we certainly missed genomic regions associated with environmental selection. More genomic data such as whole genome sequencing would be interesting to produce an analysis at finer scale. In the same way, because of data availability, we only included climatic variables in our analysis and possibly missed some other environmental factors that could have played a role in *F. sylvatica* local adaptation. However, climate is known to be the main causal factor shaping the distribution of tree species in temperate areas (Svenning and Skov 2004, Morin *et al*. 2007). Moreover, the two environmental pressures identified as drivers of the adaptation of *F. sylvatica* are congruent with the main environmental constraints driving other tree species’ distributions (e.g., *Picea glauca*, Andalo *et al*. 2005; *Picea mariana*, Beaulieu *et al*. 2004).

### Future of the species in the Alps

All general atmospheric circulation models predict that there will be major changes in temperature and rainfall by the turn of the century (IPCC 2007 and 2013). These predictions are likely to induce dramatic changes in most ecosystems around the globe and are of special concern for mountainous ecosystems (Thuiller *et al*. 2005, Thuiller *et al*. 2014, Duputié *et al*. 2015). In this context, *Fagus sylvatica* could be severely impacted (Magri 2008, Meier *et al*. 2011). According to the A1B scenario of climate change used in our simulations, the favourable zone for the survival of *F. sylvatica* in the French Alps would indeed change substantially (Fig. S6). The model predicted a drastic loss of favourable zones in lowlands and valleys and a species range shift towards higher altitudes, a pattern already observed for many plant species (Pauli *et al*. 2012, Steinbauer *et al*. 2018). This first model basically assumes a genetic homogeneity and a full dispersion of adaptive alleles. Nonetheless, we found evidence of significant co-variation between environmental and genetic gradients in the populations of beech in the French Alps. Assuming that the genetic variation identified here is indeed involved in species adaptation to climate, the species’ sensitivity to climate change could spatially vary depending on the strength of the environmental change experienced by the local populations and the availability of adaptive alleles nearby (Franks and Hoffmann 2012).

Investigating the capacity of populations’ response to climate change, we found that beech populations in high mountainous areas would need a greater change in adaptive genetic composition than would lowland or valley populations (Fig. 6A). The important changes in the climatic conditions in these areas would constrain a consequent change in the adaptive genetic component. In contrast, lowlands and valleys exhibited a weaker genetic offset but will host populations the farthest from the current locations showing the genetic composition that will be required under the future environmental conditions (geographic offset in Fig. 6B). Considering the very short time scale studied here (<100 years), we can assume that its adaptation to climate change will be mostly promoted by the genetic variation already existing between the different populations (standing genetic variation) (Barrett and Schluter 2008, Savolainen et al. 2013). Thus, any genetic adaptation would rely on either the frequency increase of alleles already present in the population or the migration of new alleles coming from locations already adapted (Jump et al. 2006, Barrett and Schluter 2008, Jump *et al*. 2009, Franks and Hoffmann 2012).

When looking more precisely at the levels of adaptive standing genetic variation in the populations, we found that some populations in the Chartreuse, Bauges and Chablais mountains currently show very low level of SGV, which would also correspond to populations where the adaptive genetic component is supposed to change the most in the next century (Fig. 6C). The small SGV found in those populations could be explained by a recent founder effect linked to the recent reforestation of the high elevation areas in the Alps. Fortunately, those populations are also strongly connected to large populations of beech in the lowland nearby from where lacking adaptive alleles could potentially migrate to balance the predicted genetic offset. In general, it seems that high mountainous populations will be able to find the needed adaptive genetic material more easily than the lowland locations, although they also show the highest need of genetic change (Fig. 6A and 6B). Such pattern is due to the geographic proximity of populations that have already adapted along the altitudinal gradient (Jump et al. 2009). In contrast, the valleys, separated from each other by high mountain ranges, could be less prompt to receive the needed adaptive genetic material. The situation is probably even more problematic for the two southernmost sampled populations (Prealpes-Azur or Verdon) where the levels of SGV are very low and the PAI very high. Those populations are already at the extreme bound of the potential genetic adaptation (high PAI) and most of the adaptive alleles are already fixed or almost fixed (low SGV), impeding their frequencies to increase even more in the future. In those circumstances, they might not be able to balance the predicted genetic offset, yet relatively weak, and track the change of climate predicted for the next decades.

By integrating together an appropriate genome scan approach, the spatially explicit estimation of future change in adaptive selective pressure and a measure of the adaptive standing genetic variation already present in the populations we have given a global picture of the potential of *Fagus sylvatica* to respond to future climate change in the French Alps. This type of integrated analyses, more and more common in the literature (Fitzpatrick and Keller 2015, Bay et al. 2018, Martins et al. 2018, Exposito-Alonso et al. 2018) is very promising to better understand the role intra-specific adaptive genetic variation can play in constraining species ability to track climate change (Jay *et al*. 2012, Alberto *et al*. 2013) and could largely benefit management strategies and conservation efforts in the future (Aitken and Whitlock 2013, Steane et al. 2014, Aitken and Bemmels 2016).

## ACKNOWLEDGEMENTS

We would like to dedicate this paper to the memory of our colleague and beloved friend Eric Bazin (1977-2017), who was involved in the origin of this work and who passed away too soon. We also would like to thank Jonas Baudry, Marion Jourdan, Marianne Bernard and Christian Miquel for their crucial help in sampling tree leaves and Etienne Klein and Sylvie Oddou-Muratorio for interesting discussions. We acknowledge support from the French National Research Agency project APPATS (ANR-15-CE02-0004) and from the project DISTIMACC (ECOFOR-2014-23, French Ministry of Ecology and Sustainable Development, French Ministry of Agriculture and Forest).

## DATA ACCESSIBILITY STATEMENT

- *Fagus sylvatica* sequences data will be available on Dryad
- Scripts and SNP dataset are available on Github: https://github.com/Capblancq/LocalAdaptationFagus

## AUTHOR CONTRIBUTIONS

E. Bazin, X. Morin and S. Lobreaux designed the study. T. Capblancq performed the analysis and treatments with the help of M. Gueguen and J. Renaud. T. C. wrote the manuscript and all authors contributed to revisions.

## SUPPORTING INFORMATION

**Table S1:** Description of sampled populations.

**Figure S1:**
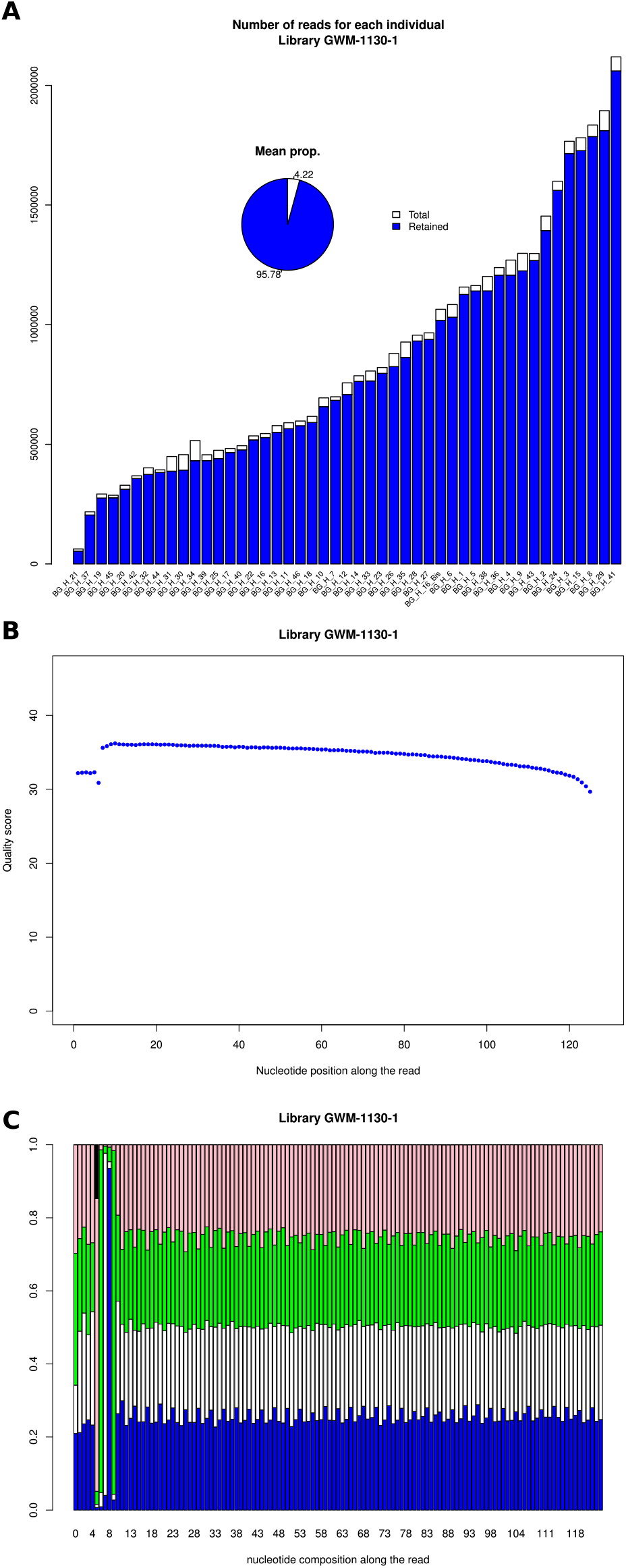
Impact of M values on polymorphism rate in ddRADseq results. M value ranges from 1 to 15.

**Figure S2:**
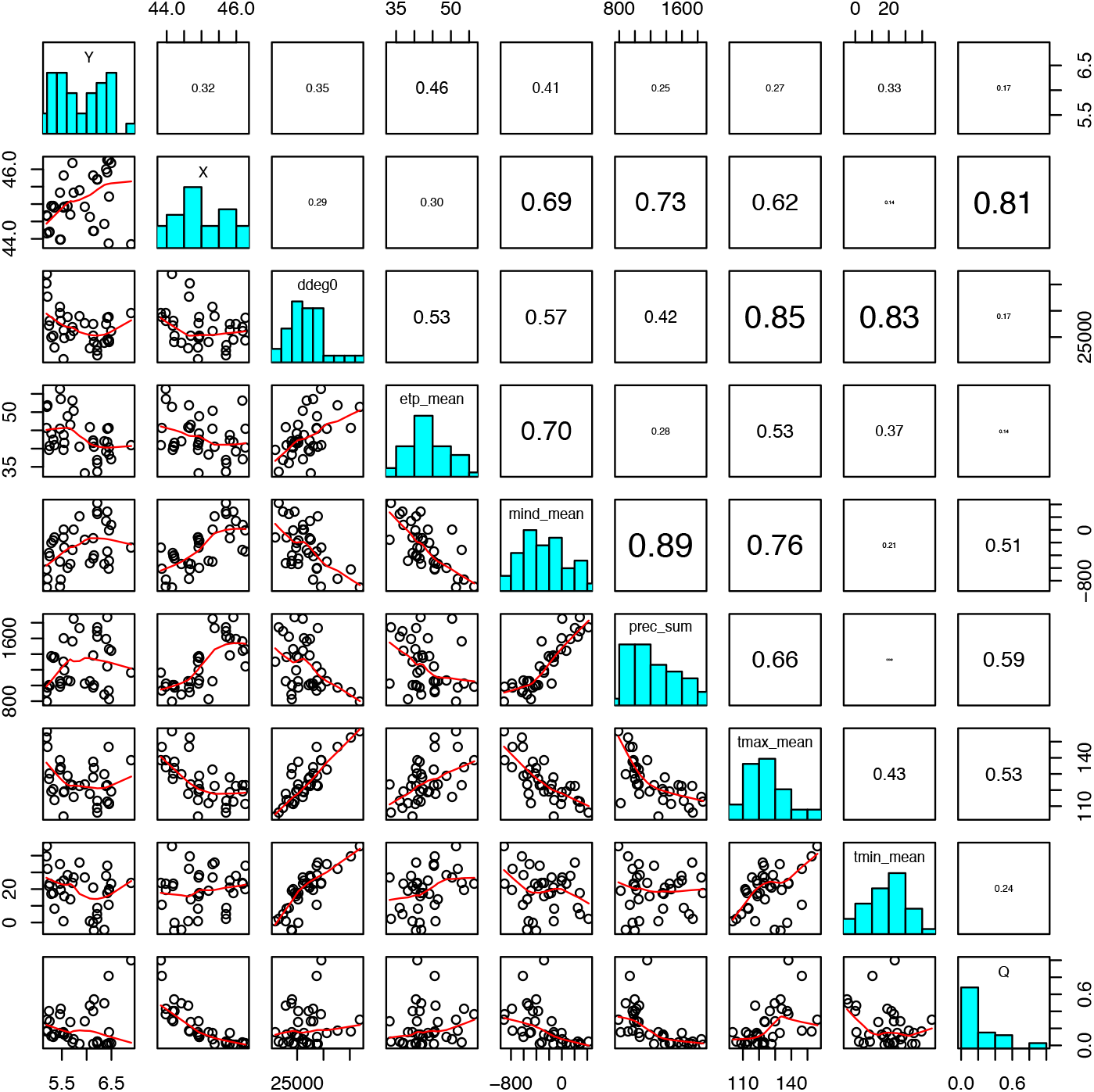
Correlations between pairs of climatic, geographic and ancestry variables. Y: latitude; X: longitude; ddeg0: the annual sum of degree-days above 0°C (degg0), the annual mean potential evapotranspiration (etp_mean), a moisture index (mind_mean), the sum of the annual precipitation (prec_sum), the maximum temperature (tmax_mean) and the minimum temperature (tmin_mean). The panels above the diagonal show the R-squared of the correlation and the panels below the diagonal show the value plot of each pair of variables.

**Figure S3:**
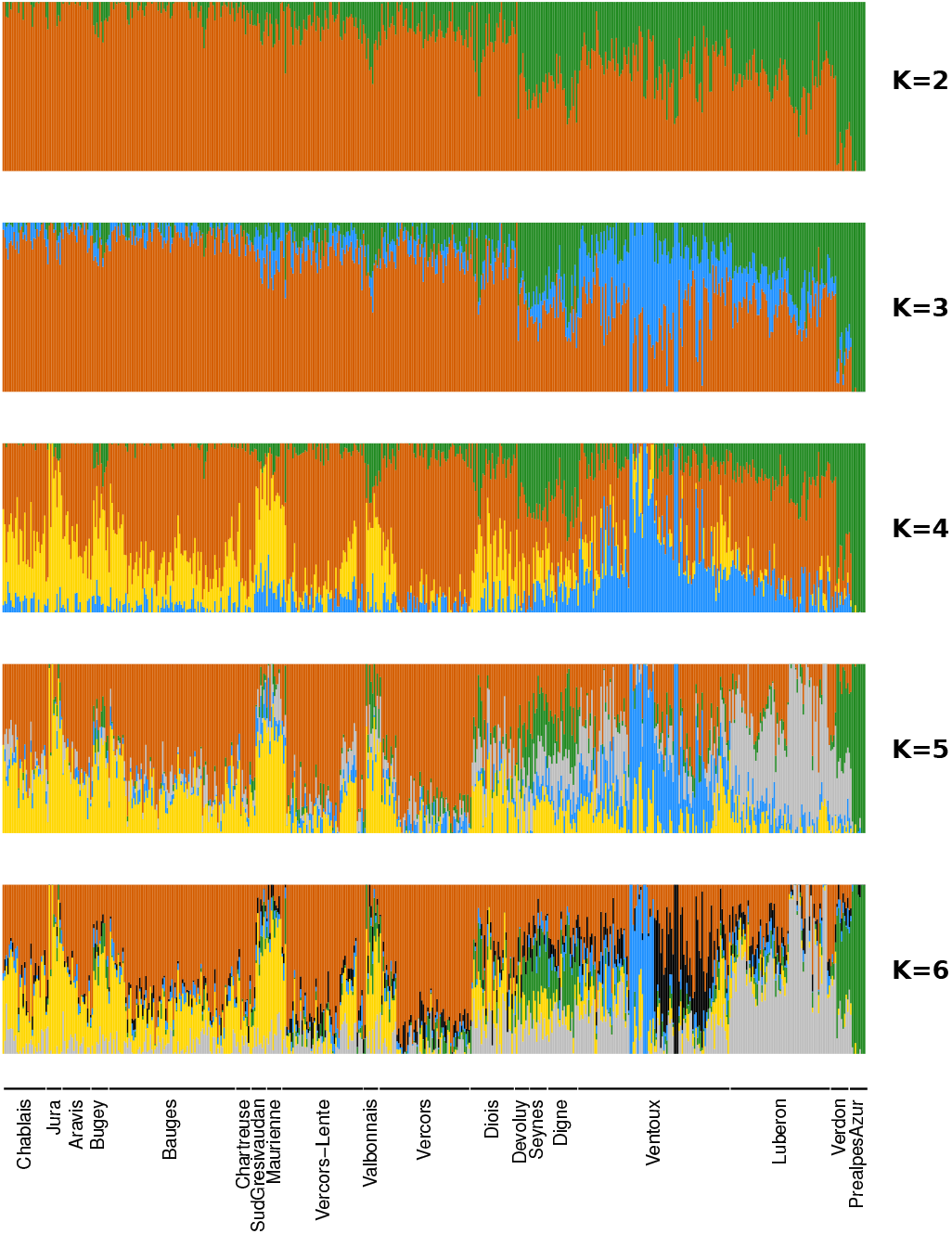
sNMF results with K ranging from 2 to 6

**Figure S4:**
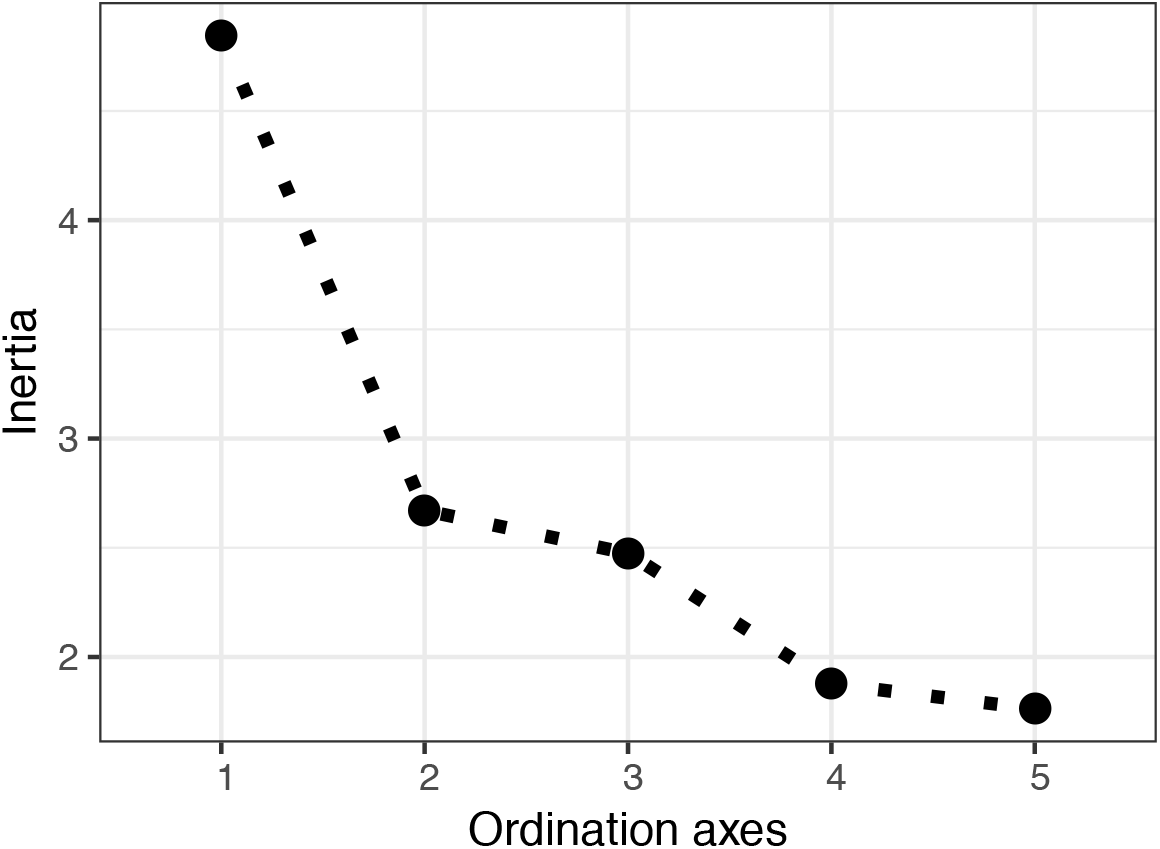
Scree plot for RDA analysis of the complete *Fagus* genetic dataset.

**Figure S5:**
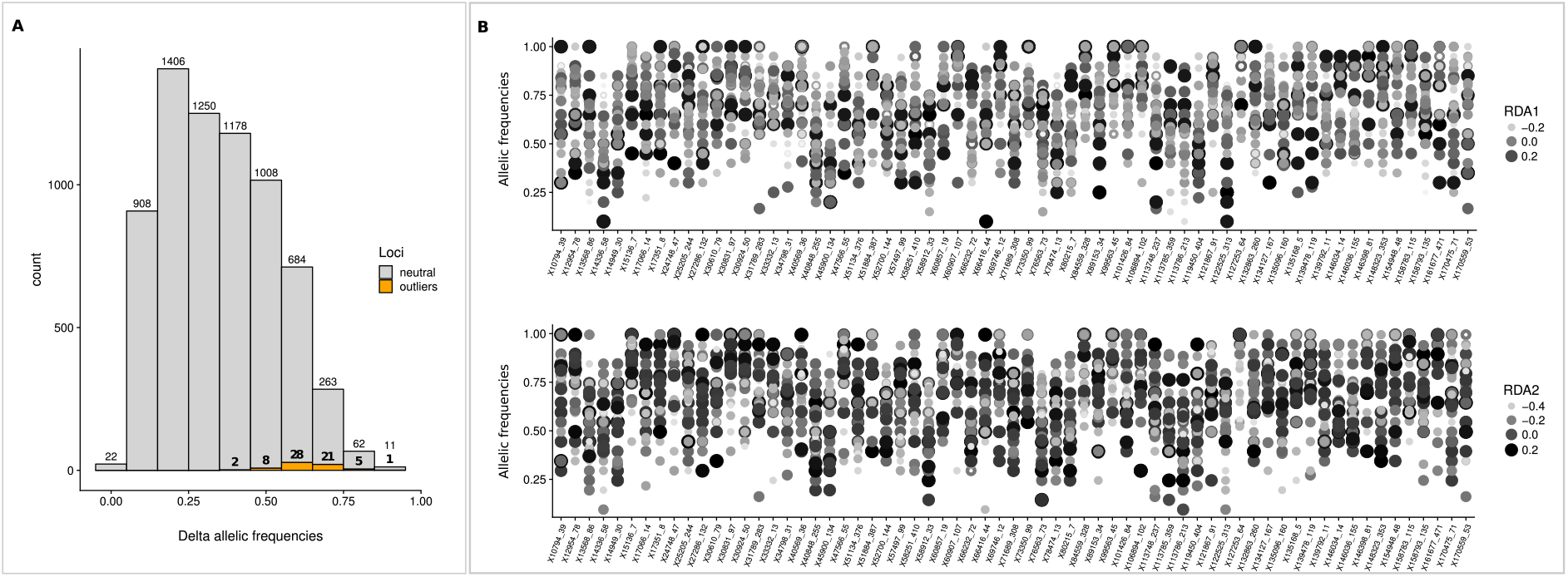
**Distribution of delta between maximum and minimum allele frequencies in sampled populations for the 4577 SNPs considered during RDA-based procedure (A).** Variation of outlier loci allelic frequencies across the sampled populations (B). The stars above the points represent the significance of the correlation between population RDA scores and allelic frequency for each locus and independently with RDA1 and RDA2 (* < 0.1; ** < 0.05; *** < 0.01).

**Figure S6:**
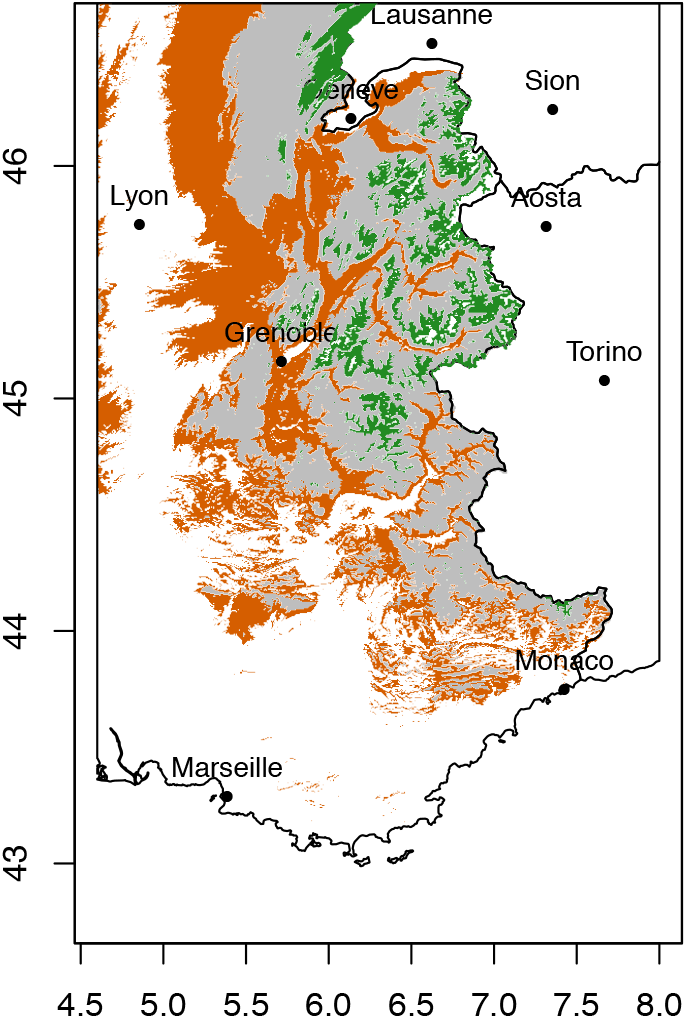
Decrease (orange) and increase (green) of favourable climatic areas for *Fagus sylvatica* in 2080. Grey pixels are favourable now and remain favourable in the future.

